# Perturbational complexity of cortical responses to thermal pain

**DOI:** 10.1101/2025.07.04.663180

**Authors:** Kora T. Montemagno, Arthur S. Courtin, Dounia Mulders, Francesca Fardo

## Abstract

While painful and non-painful thermal stimuli elicit a rich dynamical pattern of brain activity, canonical event related potentials (ERP) analyses quantify only limited aspects of this pattern. In this study, we complement the conventional ERP approach by quantifying the spatial and temporal differentiation of EEG responses to thermal stimulation using the perturbational complexity index (PCI), a complexity metric grounded in systems dynamic and information theory. Using two publicly available datasets, we computed state-transition PCI from thermal-evoked responses recorded over 32-64 scalp channels. Dataset 1 combined three stimulus intensities (10 °C, 42 °C, 60 °C) with topical application of thermosensitive TRP-channel agonists (menthol 20 %, capsaicin 1 %) or vehicle; Dataset 2 manipulated the block-wise transition probability of receiving cold (≈ 15 °C) or hot (≈ 58 °C) stimulation. PCI scaled non-linearly with temperature, being lowest at the intermediate 42 °C and highest at the cold and hot extremes (Datasets 1 and 2). PCI was sensitive both to peripheral sensitisation, as topical menthol and capsaicin selectively reduced PCI during cold stimulation (Dataset 1), and to changes in block-wise stimulus probability (Dataset 2). Across all analyses, canonical ERP peak measures (N2–P2 amplitude/latency) failed to account for PCI variance. These findings demonstrate that PCI reflects the brain’s response to exogenous, sensory-driven thermal perturbations, quantifying changes in neural complexity associated with both stimulus intensity, peripheral sensitisation and probabilistic manipulations. This supports its applicability as a measure of temporal and spatial differentiation in EEG responses relevant to pain neuroscience.

**Summary:** This study quantified the spatio-temporal complexity of electroencephalographic responses to thermal and pain stimuli. Complexity was sensitive to simulation temperature, chemical sensitization and probabilistic manipulations.

## 1 Introduction

A complex spatio-temporal pattern of brain activity accompanies the sensations elicited by both noxious and innocuous thermal stimuli. Among the most common techniques to quantify this activity, event-related potentials (ERP) allow the investigation of neural processing in the underlying network. However, conventional ERP analyses focus on parameters such as the amplitudes and latencies of specific peaks (e.g., N2–P2 complex) derived from a limited subset of electrodes, leaving the spatial dimension of the signal underexplored.

Here, we introduce a complementary analytical framework to quantify ERP dynamical richness using neurophysiological complexity-based measures. These techniques are grounded in the premise that large-scale neuronal ensembles behave as non-linear dynamical systems[17,30], and aim at characterising how information is processed across distributed brain networks[25]. Among these metrics, the Perturbational Complexity Index (PCI) provides a single numerical estimate of the spatio-temporal richness of the brain’s response to a brief perturbation. Originally introduced within the framework of Integrated Information Theory[28], the canonical protocol employs transcranial magnetic stimulation (TMS) and quantifies both the spatial/topological and temporal differentiation of the neural activity recorded with multichannel electroencephalography (EEG) using algorithmic information or recurrence-based measures[7,10,25]. While initially associated with conscious state assessment in TMS–EEG studies, PCI computation is not intrinsically tied to the mode of perturbation, so that a sensory stimulation may also be used provided that responses are reliably time-locked: we therefore investigate the feasibility and relevance of thermal stimuli as such a perturbation.

So far, a few studies have assessed sensory-evoked complexity using proxies such as entropy, functional connectivity, or algorithmic complexity. Existing work shows that complexity metrics are modulated by stimulus features and perceptual relevance across sensory modalities, including thermal and nociceptive stimulation[13,21,22,34]. Together, these findings indicate that complexity-based approaches are sensitive to both perceptual and physical properties of sensory input.

Beyond establishing thermal stimulation as a suitable perturbation for PCI, we examined whether this index tracks context-dependent modulations of pain and temperature perception. We analysed two independent EEG datasets probing distinct processing stages. In Dataset 1[12], topical application of transient receptor potential (TRP) agonists altered peripheral transduction, allowing us to assess whether PCI reflects changes in afferent gain induced by chemical sensitisation. In Dataset 2[20], block-wise manipulation of stimulus transition probabilities enabled us to test whether PCI is sensitive to contextual statistics.

We tested three related hypotheses: first, PCI would vary with stimulus intensity, with non-linear modulation emerging when stimulation becomes painful; second, chemical sensitisation would modulate this intensity–complexity relationship; and third, PCI would be sensitive to the statistics of the stimulus sequence. Together, these analyses assess PCI’s ability to measure context-dependent changes in the spatiotemporal differentiation underlying thermal and nociceptive processing.

## 2 Methods

### 2.1 Datasets description

Two publicly available datasets employing the same contact thermode (QST.Lab; Strasbourg, France) were re-analysed in this study. Ethical approval was obtained for the original studies.

Dataset 1 was originally collected by Courtin et al.[12] is available at https://dx.doi.org/10.17605/OSF.IO/VCK6S. The study investigated the effects of different TRP channel agonists on thermal event related potentials (ERP). It included 62 participants each assigned to one of four agonist groups: menthol (20%), cinnamaldehyde (10%), capsaicin (0.25%), or capsaicin (1%). Each participant was treated with an active patch (group-specific TRP agonist) on one forearm and a vehicle patch (only ethanol absolute) on the other one, in a randomised order. After the removal of each patch, the treated area was stimulated with three series of 30 thermal stimuli (target temperatures: 10°C, 42°C and 60°C; duration: 200 ms). EEG data was collected using actively shielded Ag-AgCl electrodes placed on the scalp according to the international 10/20 system (32 channels for the menthol, cinnamaldehyde and capsaicin 0.25% groups; 64 channels for the capsaicin 1% group; WaveGuard EEG cap, Advanced Neuro Technologies, Hengelo, The Netherlands).

Dataset 2 was collected by Mulders et al. [20] and is available at https://osf.io/8xvtg/. It included 31 participants who completed 10 blocks of 100 interleaved thermal stimuli delivered on the forearm. Stimuli were 250 ms long and consisted of two temperatures: I1 was set to 15°C for most participants, but could range between 15°C and 20°C; I2 was 58°C for most participants, but could be decreased to 57°C. For additional details see Mulders et al., 2023[20]. Stimuli were presented with fixed transition probabilities within each block (i.e., p(I1|I2) and p(I2|I1)). Five probability combinations were tested, each repeated twice. EEG data was recorded using 64 Ag-AgCl electrodes placed according to the international 10/10 system (WaveGuard 64-channel cap, Advanced Neuro Technologies).

### 2.2 Preprocessing

EEG recordings were pre-processed using the *mne*[15,18] and *mne_bids*[3] packages for Python (version 3.12.3). Each recording was visually inspected to identify channels that were flat or excessively noisy. Recordings with more than 30% of bad channels were excluded. Noisy channels were interpolated using spline interpolation. The EEG signal was re-referenced to the average, and ocular artifacts were identified and removed using independent component analysis (ICA). Data were filtered using a notch filter at 50Hz and a bandpass filter (0.1-45Hz, second-order Butterworth). Continuous EEG signal was then segmented into epochs from -500 ms to 1500 ms relative to stimulus onset, and baseline corrected using a window from -500 ms to -10 ms. Epochs exceeding a peak-to-peak amplitude of 200μV were excluded. For each dataset, if more than 30% of epochs were rejected within a given condition (defined as participant-patch-temperature combination for Dataset 1, or participant-block-temperature combination for Dataset 2), the entire condition was excluded from further analysis. If more than 30% of the conditions in a participant were rejected, the participant was excluded. Based on these criteria we excluded two participants in Dataset 1 and one participant in Dataset 2. Finally, we excluded one other participant in Dataset 1 who had been excluded in the original paper[12]. For all remaining participants, the EEG signal was averaged across the first 21 epochs within each condition, corresponding to the highest number of artifact-free epochs available for all conditions and participants. These condition-level averages were used in subsequent analyses.

### 2.3 Computation of perturbational complexity

The perturbational complexity index (PCI) was calculated for each condition following the state transition-based approach developed by Comolatti et al.[10]. This approach was selected over the Lempel-Ziv complexity method because it is computationally more efficient and can be applied directly in sensor space[10]. This algorithm quantifies the spatio-temporal complexity of evoked EEG responses by estimating the number of state transitions across channels and timepoints (Fig 1). First, it computes the singular value decomposition of the signal matrix to select components that account for at least 99% of the response energy (*spatial differentiation* proxy). Among these, only components with a signal to noise ratio (SNR) above a minimum threshold (*min SNR*) were retained for further analysis. On these components, complexity is estimated in terms of recurrent quantification analysis by evaluating the number of *state transitions* (NST, *temporal differentiation* proxy). For each component, voltage-amplitude differences between all time points (distance matrices) were computed separately for pre-stimulus and post-stimulus windows. These matrices were then thresholded using several levels *ε* _n_to obtain binary transition matrices indicating the number of times the signal crossed a threshold, capturing transitions between distinct states. Since the NST is quantified across several thresholds, the threshold _n_* maximizing the difference in NST between the post-and pre-stimulus windows (ΔNST) was identified and retained, and the ΔNST values were summed across components to obtain the PCI for a given condition.

**Fig 1.**
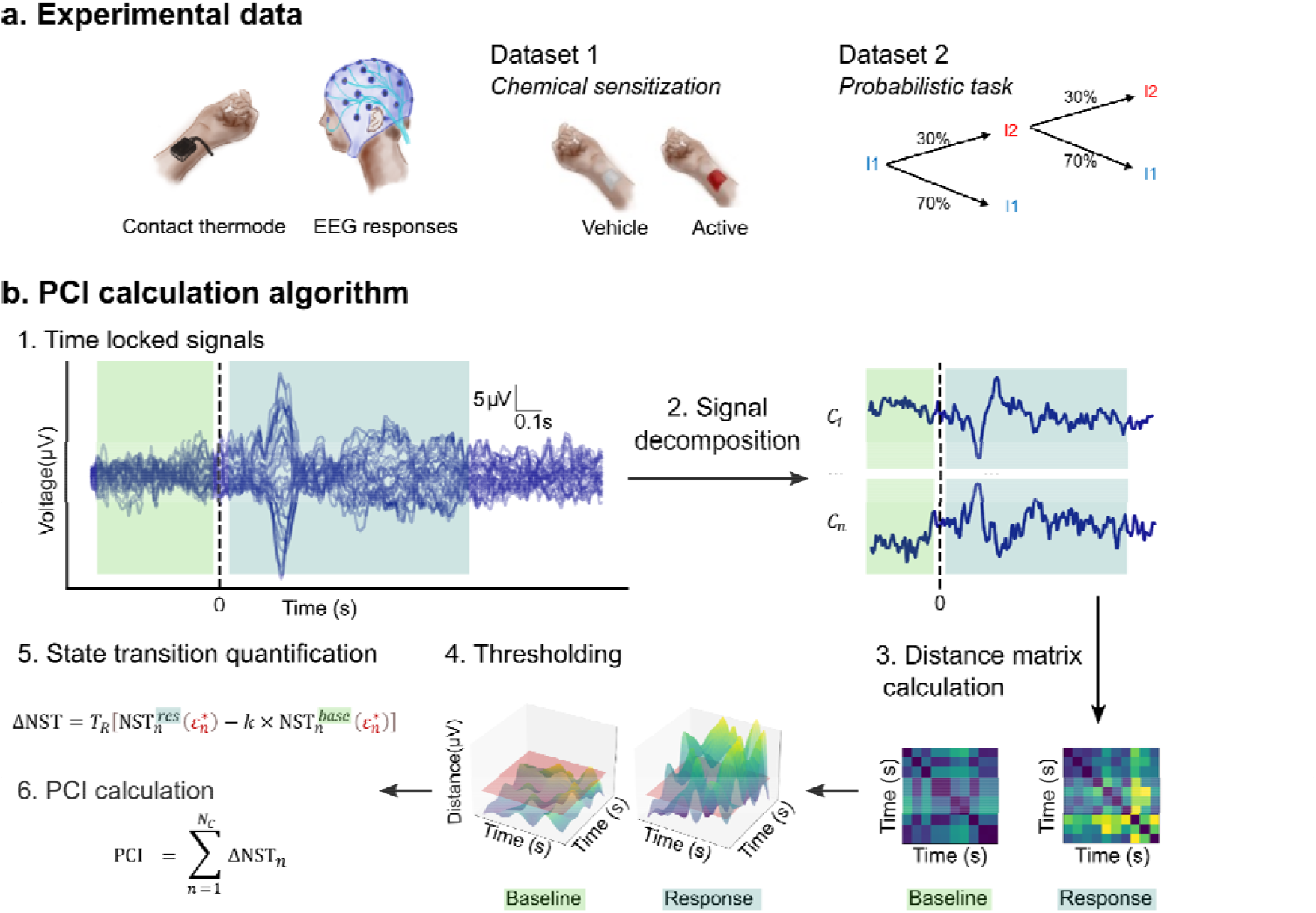
Workflow for estimating the Perturbational Complexity Index (PCI) from evoked EEG responses. PCI was estimated in two EEG datasets in which evoked responses were recorded following cold and hot stimulation delivered via a contact□thermode stimulator. Dataset 1 additionally involved the topical application of an agonist solution or vehicle (placebo□controlled), while in Dataset 2 participants completed a learning task and received sequences of cold (I1) and hot (I2) stimuli generated with five pairs of transition probabilities (here the probability of receiving a cold stimulus after a hot one p(I1|I2) = 0.7 and p(I2|I1) = 0.3). In both datasets, EEG data were identically preprocessed, and the stimulus□locked averaged signals were entered into the state□transition PCI algorithm of Comolatti□ et □al.[10], that compared the state transitions of the baseline (green) and response (light blue) time window (the dotted vertical line indicates the stimulus onset). This yielded a single PCI value for each condition (participant-□agonist□-temperature in Dataset□1; participant□-□block□-□temperature in Dataset□2). NST = number of state transitions; *ε* _n_* = threshold that maximises the difference between baseline and response NST for the component *n*; *T*_*R*_ = number of samples in the response; *k* = weight parameter between pre- and post-stimulus transitions; *N*_*c*_ = number of total components *C*.

Several considerations were applied to ensure comparability with previous work. First, components with SNR below the *min SNR* threshold were excluded. Since it is possible to expect a lower SNR for thermal evoked potentials acquired over 30-100 epochs compared to the original signal for which this algorithm was designed (200 stimuli, TMS-EEG), the *min SNR* was set to 1.1, corresponding the lowest value previously investigated[10]. Second, conditions with no surviving components (PCI = 0) were excluded, to prevent 0-inflation that could affect the linear regression models. Third, to improve comparability across conditions, PCI was computed only on the first 21 epochs within each condition, corresponding to the highest number of retained trials across all conditions in both datasets. Importantly, to ensure that the number of epochs did not significantly affect the SNR, making the index overly sensitive to noise fluctuations, we verified that the PCI values remained stable when averaging across 21, 31, 41, and 51 epochs for those conditions in Dataset 2 that included a sufficient number of trials (see Supplementary material). Fourth, while in Dataset 2 PCI was calculated using all the recording 64 channels, in Dataset 1 and in the models that grouped Dataset 1 and Dataset 2 together PCI was computed on a subsample of 32 common channels. Finally, the *k* parameter, which determines the relative weight of pre- and post-stimulus NST during thresholding, was fixed at 1.1 in accordance with the original implementation[10].

### 2.4 ERP peak parameters

To evaluate whether PCI reflects additional information compared to the amplitude or timing of canonical evoked responses, we conducted a control analysis focused on the N2–P2 complex, a well-characterised biphasic deflection elicited by thermal stimulation, typically maximal at the vertex. This analysis aimed to determine whether PCI simply reflects the magnitude of evoked activity or instead indexes distinct aspects of spatio-temporal complexity.

Following established procedures in ERP research, the N2–P2 complex was identified in the participant’s averaged signal for each condition: Fig 2 presents the grand-average ERP waveforms for each dataset and condition, showing results that are consistent with the findings reported in the original publications[12,20]. Latencies and amplitudes of the N2 and P2 components were extracted using the *mne*.*preprocessing*.*peak_finder()* function, after visual inspection of both waveforms and topography to define an appropriate time window. If peaks could not be reliably identified or if the scalp distribution was ambiguous, the corresponding condition (defined as participant–agonist–temperature in Dataset 1 or participant–block–temperature in Dataset 2) was excluded from the analysis.

**Fig 2.**
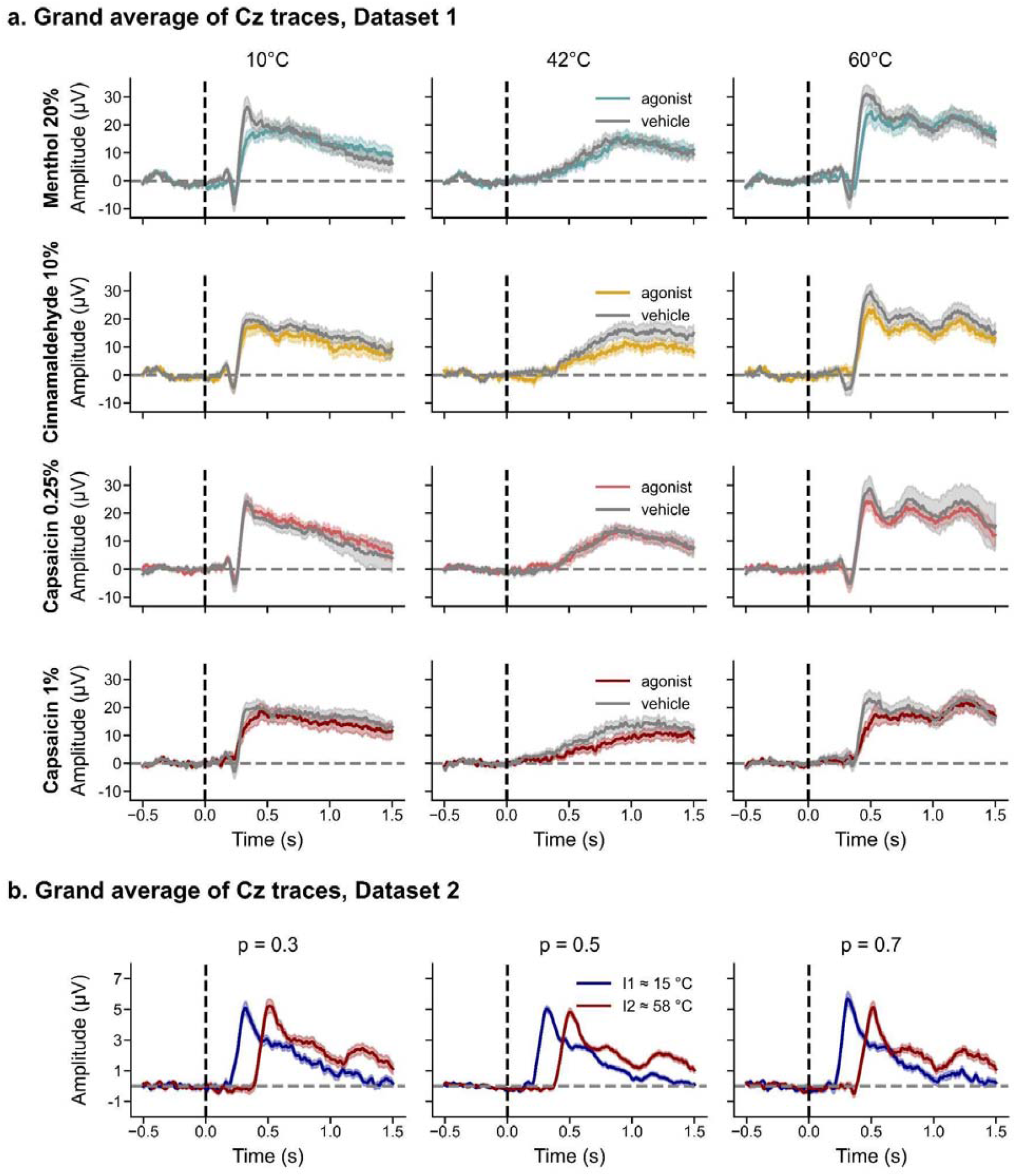
Grand-average event-related potentials (ERPs) at Cz across datasets and conditions. (a) Grand-average ERP waveforms at Cz for each combination of temperature and agonist in Dataset 1. Each row corresponds to a different agonist (Menthol 20%, Cinnamaldehyde 10%, Capsaicin 0.25%, and Capsaicin 1%), and each column shows responses to thermal stimuli at 10 °C, 42 °C, and 60 °C. Colored lines represent agonist conditions, and grey lines represent vehicle controls. Traces are referenced to the average of the mastoid channels, following the procedure used in Courtin & Mouraux[12]. (b) Grand-average ERP waveforms at Cz for Dataset 2, showing responses for three stimulus probabilities (p = 0.3, 0.5, and 0.7), and comparing low-intensity (I1 = 15 °C, blue) and high-intensity (I2 = 58 °C, red) thermal stimuli. Traces are referenced to the average of all electrodes, consistent with the approach of Mulders et al. (2023). For both datasets, solid lines indicate the mean and shaded ribbons indicate ±1 standard error of the mean (SEM). The dashed vertical line marks stimulus onset (t = 0 s). ERPs were averaged over the first 21 epochs used for PCI computation.

In Dataset 1, peak parameters were identified from the Cz electrode after averaging all valid epochs per condition. The EEG signal was re-referenced to the average of the mastoid electrodes (M1 and M2), to replicate the approach used in Courtin et al.[11]. In Dataset 2, peak extraction was also performed at Cz but on the average of the first 21 epochs per condition. Due to consistent artifact contamination at the mastoids, an average reference across all electrodes was applied to minimise condition-level exclusions. Because N2–P2 components were analysed only in relation to PCI within each dataset, and not compared across datasets, the use of different referencing schemes does not constitute a confound. This control analysis allowed us to directly compare classical ERP parameters with PCI estimates, testing whether PCI explains variance beyond canonical evoked responses.

### 2.5 Statistical analysis

All statistical analysis were performed in R (version 4.3.3[23]) using the *stats, car* and *emmeans* packages. To assess how PCI varied across conditions and datasets, we used linear regression models. The dependent variable was square-root-transformed PCI 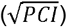 to preserve the normality assumption. Distribution of the residuals was assessed using quantile-quantile (QQ) plots to verify test assumptions. If a data point strongly violated normality, it was removed, and the model was re-estimated (2/366 data points excluded for Model 1, 1/630 for Model 2, 3/366 for Model 3, 2/572 for Model 4 Dataset 1 cold, none for the other models). When post-hoc comparisons were required, p-values were adjusted using the Holm-Bonferroni procedure to control for the family-wise error rate (α = .05). We constructed four models to assess condition-specific effects and potential covariates.

#### 2.5.1 Dataset 1: Effects of temperature and TRP agonists on PCI

To test the effect of different TRP channel agonists on PCI across three temperatures (10°C, 42°C, 60°C), we fitted the following model:

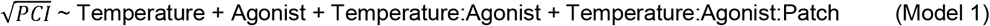

where Temperature, Agonist, and Patch were categorical predictors, with Patch indexing whether stimulation was preceded by an active or vehicle treatment. Since this model employs only categorical variables, we used a Type III ANOVA to investigate the significance of effects.

#### 2.5.2 Dataset 2: Effects of temperature and probabilistic context on PCI

To assess the effect of the absolute probability of receiving a cold stimulus (*p*(I1) = 1 - *p*(I2)) on PCI values across different blocks, we tested the following model:

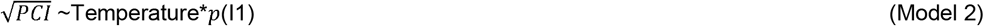

where Temperature was a categorical predictor, (reference = I1), and *p*(I1) a metric variable, mean-centered prior to the regression. The symbol * indicates that both the single variables and their interaction were considered (i.e. Temperature + *p*(I1) + Temperature: *p*(I1)). Regression coefficients and marginal effects were employed to evaluate the significance of effects.

#### 2.5.3 Comparison across datasets

To assess the effect of Temperature across the 2 datasets, we tested the following model:

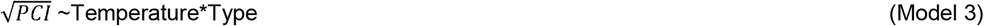

where Temperature was a metric variable encompassing all the temperatures in the two datasets, and Type represents the categorical variable encoding the stimulus type (cold vs hot). As in model 2, regression coefficients and marginal effects were employed to evaluate the significance of effects.

#### 2.5.4 ERP control analysis

To test whether PCI could be explained by canonical evoked responses, we modelled PCI as a function of N2 and P2 latency and amplitude:

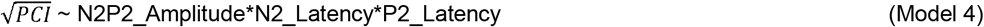

Predictor variables were mean-centred and scaled to reduce multicollinearity. Regression coefficients and marginal effects were employed to evaluate the significance of effects. The 95% confidence interval for the model’s R^2^ was estimated using the *Ci.rsq()* function from the *psychometric* package in R.

## 3 Results

### 3.1 PCI varies as a function of temperature

To assess whether PCI reflects differences in stimulus intensity, we first examined the effect of temperature on PCI for each dataset separately. In Dataset 1, participants received three temperature stimuli (10°C, 42°C, and 60°C). Model 1 revealed a significant main effect of temperature on 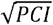 (F(2,320) = 16.58, *p* < .001). Pairwise comparisons within the vehicle condition indicated that 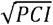 at 42°C was significantly lower than at both 10°C and 60°C (difference Δ42-10 = -1.43, SE_42-10_ = 0.13, *p* < .001; Δ_42-60_= -1.21, SE_42-60_ = 0.13, *p* < .001).

In Dataset 2, participants received two temperature stimuli (I1≈15°C, I2≈58°C). Model 2 revealed a significant effect of Temperature, with lower values associated with hot stimuli (average marginal effect AME of Temperature I2: = -0.35, SE = 0.09, *p* < .001).

Model 3 assessed the main effect of Temperature on both datasets, allowing us to control for stimulus type (Type = hot or cold) and to investigate how PCI varies across all the considered temperatures. The results of the regression showed significant effects of both Temperature (regression coefficient β = -0.10, SE = 0.04 *p* = .014), Type (β_hot_ = -5.68, SE = 0.72, *p* < .001) and the interaction term Type × Temperature (β = 0.17, SE = 0.04, *p* < .001), indicating that stimulus type influences the Temperature slope. Indeed, the marginal effect of temperature varied for the cold and hot stimuli (Marginal Effect ME_cold_ = -0.10, SE = 0.04, *p* =.014; ME_hot_ = 0.07, SE = 0.01, *p* < .001), suggesting a U-shaped relationship: the more the stimulus temperature departs from the baseline, the more the complexity increases.

Together, these findings indicate that PCI is modulated by stimulus temperature, with increased complexity observed at temperatures further from baseline (Fig 3), and a slightly more pronounced effect within the cold temperature range. This effect may be due to the limited range of temperatures tested, their asymmetry in terms of distance from baseline, differences in perceived intensity, or by distinct neural processing engaged by cold and hot stimuli.

**Fig 3.**
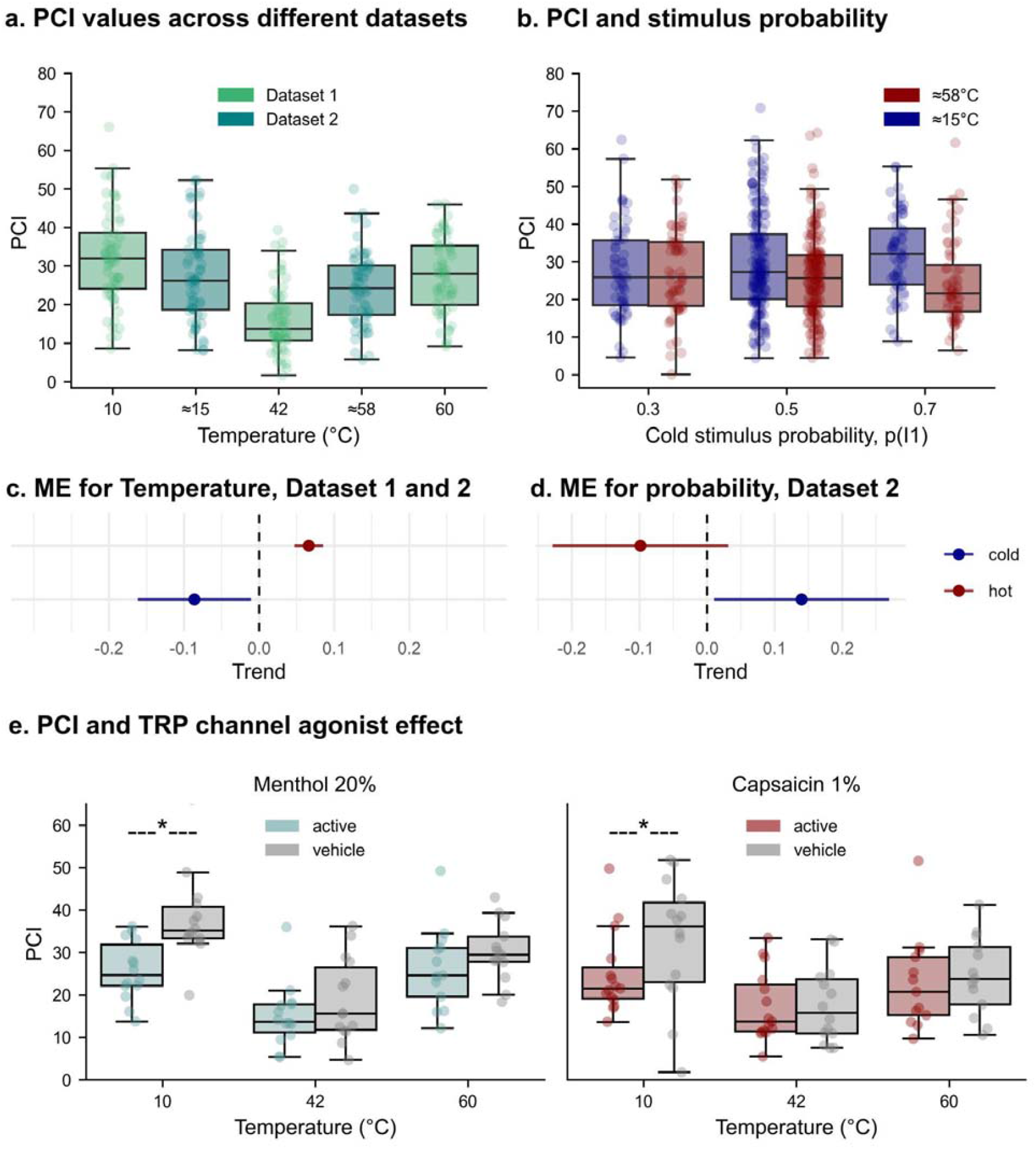
Results from linear regression models. **a**. Box□and□whisker plots summarise PCI values computed on the 32□channel montage for each temperature in Dataset □1 (Courtin & Mouraux[12]; 10□°C, 42□°C, 60□°C, patch=“vehicle”) and Dataset □2[20]; I1≈15□°C and I2≈58□°C, transition probabilities = 0.5). The PCI varied non-linearly with temperature across both datasets, reaching highest values at the temperature extremes (10□°C and 60□°C, Dataset 1) and a minimum at the intermediate 42□°C. **b**. Box-and-whisker plots show raw PCI scores calculated from the full 64-channel montage, grouped by the block-wise probability of receiving a cold stimulus I1 ≈□15□°C (0.3,□0.5,□0.7). Colours distinguish the two temperatures: hot I2 ≈□58□°C (red) and cold I1 ≈□15□°C (blue). Consistent with the statistical analysis (Sections 3.1 and 3.3), it is possible to observe a Temperature effect on PCI, being it lower for hot stimuli (AME = - 0.35, p < .001). Furthermore, complexity increases when *p*(I1) increases for cold stimuli (ME_cold_ = 0.140, *p* = .034). For visualisation of these statistical results, figure **c**. and **d**. show a point-and-whisker plot depicting the marginal effects related to temperature (cold vs hot, model 3), and to the absolute probability of receiving a stimulus, respectively. **e**. Box□and□whisker plots show raw PCI scores (32□channel montage) for menthol□20□% (left) and capsaicin□1□% (right). For each agonist, participants received a vehicle patch and an active patch before stimulation at 10□°C, 42□°C, and 60□°C; the patch order was counter□balanced within subjects. Both agonists selectively lowered complexity at 10□°C, whereas no consistent patch effect emerged at 42□°C or 60□°C. The significant differences between agonist and vehicle conditions for each temperature are indicated in the plot by the symbol *. For the other agonists, see Supplementary materials. In each Box-and-whisker plot, the single PCI estimate for each condition is shown as a semi□transparent dot. Boxes denote the median and inter□quartile range (IQR), whiskers extend to 1.5□×□IQR.

**Fig. 4.**
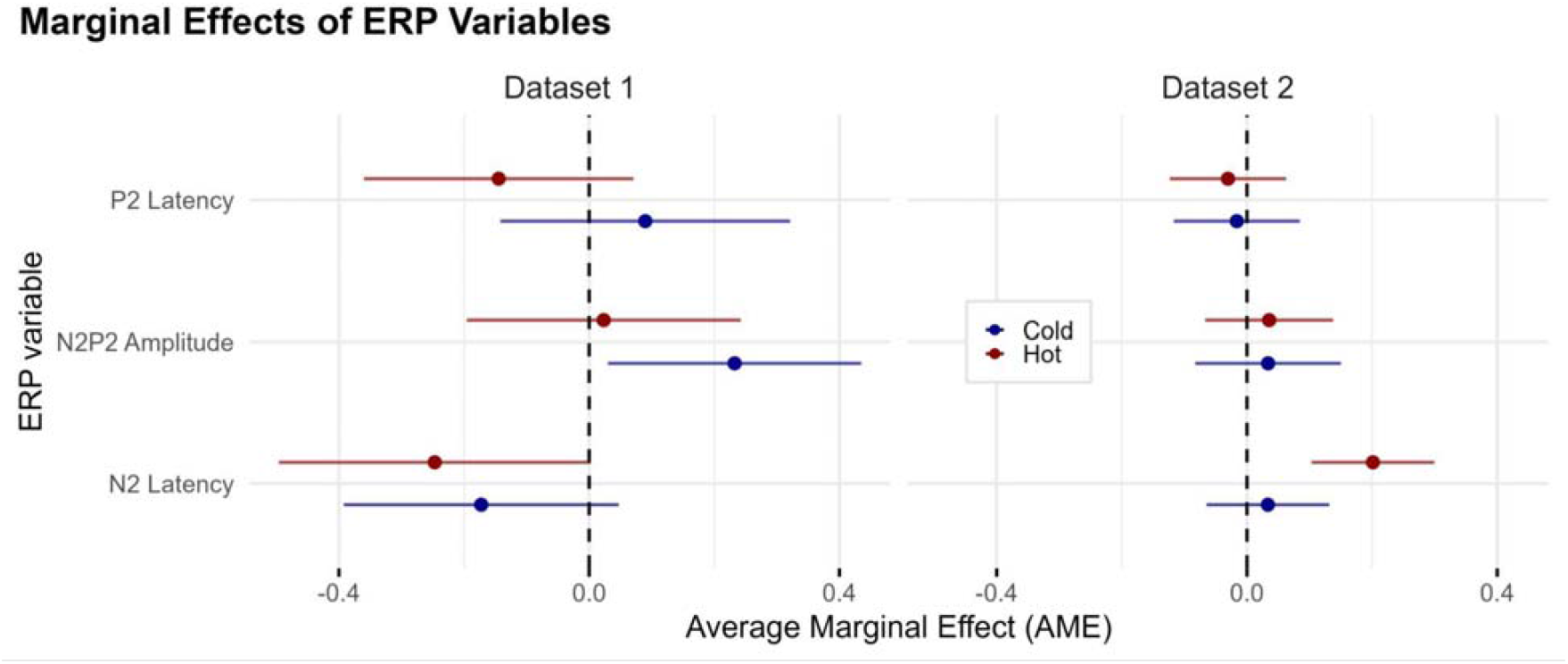
Relationship between PCI and ERP parameters. Point-and-whisker plots for each dataset and each stimulus type (cold and hot) depicting the marginal effect of each ERP parameter on PCI. Whiskers extend to 95% confidence interval. No statistically significant effect was found for most ERP parameters in both datasets, with the exception of N2P2 amplitude effect on cold stimuli PCI, Dataset 1 (AME = 0.23, SE = 0.10, *p* = .024) and N2 latency on hot stimuli PCI in Dataset 2 (AME = 0.20, SE = 0.05, *p* < .001).

### 3.2 PCI varies as a function of TRP agonist application

Dataset 1 included within-subject comparisons of electrophysiological responses following application of either a TRP agonist or vehicle patch to the forearm. Using Model 1, we found that the Temperature × Agonist × Patch interaction was significant (F(12,320) = 2.24, *p* = .010), prompting further analysis within each patch condition. Notably, Agonist effect was significant, but Temperature x Agonist effects was not not significant (F(3,320 = 3.08, *p* = .028; F(6,320) = 1.24, *p* = .286, respectively), suggesting that, while some baseline differences between the groups may be present, there may be an effect specific to the three-way interaction.

Significant decreases in 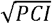 were observed under cold stimulation (10°C) for both capsaicin 1% (Δ_vehicle-agonist_= 0.91, SE_vehicle-agonist_= 0.37, *p* = .040) and menthol 20% (Δ_vehicle-agonist_= 1.23, SE_vehicle-agonist_= 0.35, *p* = .002). Agonist manipulation did not lead to significant differences at 42□°C or 60□°C (Fig 3), and no effects were observed on any temperature under capsaicin .25% or Cinnamic aldehyde 10%.

Overall, topical capsaicin and menthol selectively reduced complexity during cold stimulation, with no detectable influence at warmer temperatures.

### 3.3 PCI is affected by probabilistic manipulations

Dataset 2 allowed us to examine whether cortical complexity is modulated by expectation as, across successive blocks, the absolute probability of receiving a cold (I1≈15°C) stimulus was set to *p*(I1) = 0.3, 0.5,□or 0.7. Notably, the probability of receiving a hot (I2) stimulus varied accordingly but in the opposite fashion, being *p*(I2) = 1 - *p*(I1). Thus, three different combinations of absolute stimulus probabilities were present in the dataset.

Model 2 assessed the main effect of this stimulus probability change and its interaction with stimulus temperature on 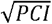. The regression coefficients associated with *p*(I1) (β = 0.14, SE = 0.07, *p* = .034) and the interaction term *p*(I1) x Temperature (β = -0.24, SE = 0.09, *p* = .011) were significantly different from zero. When fixing the Temperature to I1 (cold), *p*(I1) at I1 temperature marginal effect was significant and positive (ME_cold_ = 0.14, SE = 0.07, *p* = .034), indicating that complexity increased with the absolute probability of cold stimuli. In contrast, for I2 (hot) stimuli, the effect was negative and not significant (ME_hot_ = - 0.10, SE = 0.07, *p* = .140). Since *p*(I2) = 1 - *p*(I1), this implies that complexity tended to increase with the absolute probability of a given stimulus, more reliably so for cold temperatures.

### 3.4 PCI is largely independent of conventional N2–P2 peak metrics

To determine whether PCI merely tracks the latency or amplitude of the N2–P2 complex, we fitted Model 4 separately for each dataset and temperature. In Dataset 1, for cold stimuli, the marginal effect of N2-P2 amplitude on 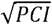 was significant (AME = 0.23, SE = 0.10, *p* = .024), but no significant effect was found for N2 and P2 latency (AME = -0.17, SE = 0.11, *p* = .124; AME = 0.09, SE = 0.12, *p* = .448, respectively). For hot stimuli, no marginal effect proved significant, even though a negative trend was present for N2 latency (N2 latency: AME -0.25, SE = 0.13, *p* = .052; P2 Latency: AME = -0.14, SE = 0.11, *p* = .188; N2-P2 Amplitude: AME = 0.02, SE = 0.11, *p* = .835).

In Dataset 2, no significant marginal effects were found for all the regressors for cold stimuli (N2 latency: AME 0.03, SE = 0.05, *p* = .504; P2 Latency: AME = -0.02, SE = 0.05, *p* = .753; N2-P2 Amplitude: AME = 0.03, SE = 0.06, *p* = .570). For hot stimuli, only N2 latency had a significant effect on complexity (N2 Latency: AME = 0.20, SE = 0.05, *p* < .001, P2 Latency: AME = -0.03, SE = 0.05, *p* = .523; N2-P2 Amplitude: AME = 0.04, SE = 0.05, *p* = .496).

To summarise, these results, while showing some sporadic association between peak parameters and complexity, do not point towards a consistent effect, suggesting that PCI may explain other features associated with EEG responses to thermal stimuli. Furthermore, the models explained little variance: R^2^ never exceeded □.10 and every 95□% confidence interval included zero for both cold (Dataset□1: R^2^□=□.10, CI□[□.22,□.42]; Dataset□2: R^2^□=□.02, CI□[□.20,□.25]) and hot stimuli (Dataset 1, *R*^2^= .10, CI [-.27;.48]; Dataset 2, *R*^2^= .05, CI [-.23;.34]).

These results suggest that classical ERP peak parameters account for only a small fraction of PCI variability, reinforcing the idea that PCI quantifies aspects of spatio-temporal complexity beyond traditional amplitude□latency measures.

## 4 Discussion

The present study used sensory□evoked perturbational complexity to characterise how the human cortex integrates and differentiates information under graded thermal input, peripheral sensitisation, and probabilistic manipulation, establishing that complexity analyses quantify objective and subjective stimulus characteristics.

Four key findings emerged: (1) cortical complexity scaled with stimulus intensity in a non-linear manner; (2) peripheral sensitisation reduced PCI during cold perception; (3) probabilistic manipulations have a detectable impact on PCI; and (4) PCI captured aspects of the evoked response beyond those reflected in conventional ERP measures of latency and amplitude.

In response to graded thermal input, PCI systematically varied with the stimulus distance from baseline skin temperature. In Dataset 1, PCI exhibited a U-shaped pattern, reaching its minimum at 42□°C and increasing significantly at both 10□°C and 60□°C. Notably, Notably, in Dataset 2 cold stimuli elicited slightly higher PCI values than hot ones (≈15°C vs ≈58°C), suggesting that intense cold may recruit a more diverse or distributed set of cortical states than intense hot. This non-linear profile aligns with previous findings, which show that thermal extremes yield a decrease in EEG alpha power, likely reducing low-frequency synchronisation[12,27]. The differential effects of cold and hot stimulation may emerge from differences in information processing, as suggested by the difference in spatio-temporal and spectral characteristics of brain activity elicited by cold and hot stimuli[32]. However, such effects may also stem from imperfect matching of stimuli across modalities, for example in terms of perceived intensity or painfulness.

Another key finding is that PCI was sensitive to peripheral sensitisation induced by TRP agonists. In Dataset 1, topical application of menthol (20%) and capsaicin (1%) significantly reduced PCI in response to cold stimulation (10□°C), relative to vehicle patches. This reduction in PCI may suggest that complexity is dependent on the state of the system prior to the stimulus, perhaps more than on the differences in afferent gain. Indeed, menthol may reduce perceived intensity of cold in some contexts[1] but capsaicin typically does not[12], despite both increasing spontaneous sensations during patch application. PCI may thus reflect not only the properties of the stimulus, but also the brain’s readiness to process it in a flexible, integrative manner, highlighting that spatio-temporal complexity may exhibit some degree of state dependency[5].

These results are consistent with prior ERP findings from the same dataset, where menthol application led to reduced N2–P2 amplitudes and prolonged latencies, and capsaicin similarly reduced amplitude without significantly affecting latency[12]. Critically, even though in the original paper significant variations in amplitude and latencies could be observed also for hot-evoked responses in response to menthol 20% and capsaicin 1% patches, such effects were not significant for PCI. Similarly, the application of cinnamic aldehyde 10% did not lead to significant changes in complexity, despite having been reported to influence amplitudes and latencies for heat-evoked responses. These discrepancies could reflect that, while classical ERPs capture signal magnitude and timing at specific scalp locations and may be correlated to some extent to spatio-temporal complexity, PCI offers a global, state-based account of how stimulus-evoked activity unfolds across time and space, possibly providing different information. PCI reductions cannot indeed be fully explained by changes in conventional ERP features. While decreases in N2–P2 amplitude or shifts in latency could hypothetically lower the number of high-SNR components retained during the PCI calculation leading to a PCI decrease, regression analyses showed that ERP parameters explained only a small fraction of PCI variance (R^2^ ≤ .10 across conditions).

Finally, when assessing PCI sensitivity to probabilistic manipulations (i.e., sequence statistics) we found evidence of modulation of PCI by the probability of receiving a cold stimulus in Dataset 2. Concurrently, existing evidence points towards an effect of sequence statistic on complexity, as studies using laser-evoked potentials have shown that the N2-P2 waveform can be modulated by stimulus novelty in combination with saliency, unpredictability, and behavioural relevance[24,29]. However, whether our findings genuinely reflect changes in cognitive processing related to the probabilistic manipulation is not certain. Because stimulus probabilities were asymmetrical in some blocks and we analysed the first 21 artefact-free epochs for each temperature-probability combination, stimuli with a low probability were on average delivered later in the sequence than stimuli with a high probability. Thus, we could not control for the effect of habituation across trials, which is known to influence brain responses to thermal stimuli and may have lowered the complexity associated with low-probability conditions[16].

### Theoretical implications: a complexity framework for pain and temperature perception

PCI was originally developed within the framework of Integrated Information Theory to quantify the brain’s capacity for consciousness based on the complexity of cortical responses to direct perturbation, typically via transcranial magnetic stimulation (TMS)[7,10,28]. It reflects the degree to which neural activity is both *differentiated* (i.e., rich in diverse, time-varying components) and *integrated* (i.e., distributed across interconnected sources).

In this framework, *differentiation* corresponds to the number and diversity of unique neural responses: the more diverse the unique neural responses, the more complex and unpredictable the signal will be over time[19]. This notion is closely linked to *temporal differentiation*, as the recorded signal reflects the composition of multiple irreducible time series. In contrast, *integration* reflects the coherent propagation of activity across brain regions, enabling distant neural populations not directly affected by the perturbation to respond in a coordinated manner[19]. This is conceptually related to *spatial differentiation*, since functional integration requires interconnections among spatially distributed sources. Notably, there is an overlap between IIT constructs of *differentiation* and *integration* and the measurable signal properties of *temporal* and *spatial differentiation*. For example, the diversity of unique neural responses (*IIT differentiation*) depends in part on the variety of contributing brain regions (*spatial differentiation*), while the coordinated propagation of activity (*IIT integration*) shapes the structure of the temporal signal (*temporal differentiation*)[25]. PCI concurrently captures both dimensions simultaneously[7]. Indeed, TMS-evoked PCI has been shown to clearly distinguish between states of consciousness, suggesting that this combined approach more accurately reflects the richness of brain dynamics[6,8,26,31].

Extending this framework to sensory-evoked PCI associated with painful and non-painful thermal stimuli introduces a different interpretive domain. Unlike transcranial perturbations to probe consciousness states, thermal stimulation modulates the contents of ongoing conscious perceptual experience. Yet, the same principles apply: high PCI may indicate that the stimulus engages brain-wide, temporally rich processing that supports adaptive, flexible responses; low PCI may signal reduced integration or differentiation, consistent with less flexible or stereotyped neural responses. From this perspective, PCI quantifies properties of the distributed network dynamics contributing to the construction of thermal and painful experiences, rather than the presence or absence of consciousness per se, making it an interesting measure for future work on state-dependent modulation of pain processing.

Within predictive coding frameworks[14], a low complexity could correspond to failures in hierarchical inference, such as maladaptive precision weighting or impaired model updating, both of which have been implicated in chronic pain pathophysiology[9]. Notably, this interpretation aligns with evidence from chronic pain research, where states of increased sensitivity or persistent pain are associated with less variable, more spatially and temporally constrained cortical dynamics[2,4,33]. Therefore, future research may investigate PCI in response to thermal stimuli delivered to individuals with chronic pain.

Further research should systematically investigate how PCI varies within a single individual over time, across stimulation modalities, recording techniques (e.g., magnetoencephalography), and contextual modulation to fully characterise its utility as a systems-level marker of adaptive or maladaptive brain states. Nonetheless, these findings open new avenues for the application of complexity-based metrics in pain and temperature neuroscience, particularly in understanding how endogenous state changes and exogenous manipulations jointly shape the brain’s capacity to process and integrate nociceptive information.

## Conclusion

Perturbational complexity has primarily been applied as a global index of consciousness, showing that reduced complexity reflects loss of integration and differentiation of the information carried on by a TMS perturbation. Here, we extended the use of PCI to sensory-evoked perturbations, applying it to characterise the network states underlying thermal and nociceptive processing. Across two independent datasets, we found that PCI was sensitive to both stimulus intensity, peripheral sensitisation via TRP agonists, and probabilistic manipulations. The negligible association between PCI values and conventional ERP parameters highlights PCI capacity to quantify large-scale changes in the spatio-temporal organisation of evoked cortical responses.

By assessing PCI sensitivity to stimulus features and contextual modulation, our findings support the use of perturbational complexity-based metrics as a sensitive tool for probing the large-scale neural dynamics associated with thermosensation and pain perception.

## 5 Acknowledgments

Generative AI tools, specifically ChatGPT, were utilised in this research to assist in text and code review (i.e. grammar correction, error debugging) and generation of non critical code (i.e. plot styling, checks, automatisation). The AI outputs have been rigorously reviewed and verified for accuracy, and their use has been disclosed in compliance with Aarhus University guidelines. The researchers take full responsibility for the content of the published article.

## Data and Code Availability

All data are publicly available via the Open Science Framework

(Dataset 1: https://dx.doi.org/10.17605/OSF.IO/VCK6S; Dataset 2: https://osf.io/8xvtg/). The analysis code used in this study is available at: https://github.com/Body-Pain-Perception-Lab/pain-PCI_pipeline/tree/main/publication.

## Authors contributions

KTM: Conceptualisation, data curation, formal analysis, funding acquisition, methodology, project administration, software, validation, visualisation, original draft, writing – review & editing.

ASC: Investigation, methodology, resources, supervision, validation, original draft, writing – review & editing.

DM: Investigation, methodology, resources, writing – review & editing.

FF: Funding acquisition, methodology, project administration, supervision, original draft, writing – review & editing.

## Funding

This work was supported by a Neuroscience Academy Denmark (NAD) fellowship (KTM, Lundbeck Foundation grant R389-2021-1596, Neuroscience Academy Denmark); the European Research Council (ERC-2020-StG-948838; FF and ASC) and the Lundbeck Foundation (R436-2023-991; FF).

## Declaration of Competing Interest

Authors report no conflicts of interest.

## Notes

https://dx.doi.org/10.17605/OSF.IO/VCK6S

https://osf.io/8xvtg/

https://github.com/Body-Pain-Perception-Lab/pain-PCI_pipeline/tree/main/publication

### Competing Interest Statement

The authors have declared no competing interest.

### Summary of Updates

Introduction and discussion have been re written to make the manuscript more pertinent to the pein field. Figures have been improved for clarity. A small change in the data analysis of model 1 (outlier removal) has been done to ensure consistency between the models

## References

[1] Andersen HH, Poulsen JN, Uchida Y, Nikbakht A, Arendt-Nielsen L, Gazerani P. Cold and L-menthol-induced sensitization in healthy volunteers--a cold hypersensitivity analogue to the heat/capsaicin model. Pain 2015;156:880–889.

[2] Apkarian VA, Hashmi JA, Baliki MN. Pain and the brain: Specificity and plasticity of the brain in clinical chronic pain. PAIN 2011;152:S49.

[3] Appelhoff S, Sanderson M, Brooks TL, Vliet M van, Quentin R, Holdgraf C, Chaumon M, Mikulan E, Tavabi K, Höchenberger R, Welke D, Brunner C, Rockhill AP, Larson E, Gramfort A, Jas M. MNE-BIDS: Organizing electrophysiological data into the BIDS format and facilitating their analysis. J Open Source Softw 2019;4:1896.

[4] Baliki MN, Geha PY, Apkarian AV, Chialvo DR. Beyond Feeling: Chronic Pain Hurts the Brain, Disrupting the Default-Mode Network Dynamics. J Neurosci 2008;28:1398–1403.

[5] Battaglia D. Function Follows Dynamics: State-Dependency of Directed Functional Influences. In: Wibral M, Vicente R, Lizier JT, editors. Directed Information Measures in Neuroscience. Berlin, Heidelberg: Springer, 2014. pp. 111–135. doi:10.1007/978-3-642-54474-3_5.

[6] Breyton M, Fousek J, Rabuffo G, Sorrentino P, Kusch L, Massimini M, Petkoski S, Jirsa V. Spatiotemporal brain complexity quantifies consciousness outside of perturbation paradigms. eLife 2025;13. doi:10.7554/eLife.98920.2.

[7] Casali AG, Gosseries O, Rosanova M, Boly M, Sarasso S, Casali KR, Casarotto S, Bruno M-A, Laureys S, Tononi G, Massimini M. A Theoretically Based Index of Consciousness Independent of Sensory Processing and Behavior. Sci Transl Med 2013;5:198ra105–198ra105.

[8] Casarotto S, Comanducci A, Rosanova M, Sarasso S, Fecchio M, Napolitani M, Pigorini A, G. Casali A, Trimarchi PD, Boly M, Gosseries O, Bodart O, Curto F, Landi C, Mariotti M, Devalle G, Laureys S, Tononi G, Massimini M. Stratification of unresponsive patients by an independently validated index of brain complexity. Ann Neurol 2016;80:718–729.

[9] Chen ZS, Wang J. Pain, from perception to action: A computational perspective. iScience 2023;26:105707.

[10] Comolatti R, Pigorini A, Casarotto S, Fecchio M, Faria G, Sarasso S, Rosanova M, Gosseries O, Boly M, Bodart O, Ledoux D, Brichant J-F, Nobili L, Laureys S, Tononi G, Massimini M, Casali AG. A fast and general method to empirically estimate the complexity of brain responses to transcranial and intracranial stimulations. Brain Stimulat 2019;12:1280–1289.

[11] Courtin AS, Maldonado Slootjes S, Caty G, Hermans MP, Plaghki L, Mouraux A. Assessing thermal sensitivity using transient heat and cold stimuli combined with a Bayesian adaptive method in a clinical setting: A proof of concept study. Eur J Pain 2020;24:1812–1821.

[12] Courtin AS, Mouraux A. Combining Topical Agonists With the Recording of Event-Related Brain Potentials to Probe the Functional Involvement of TRPM8, TRPA1 and TRPV1 in Heat and Cold Transduction in the Human Skin. J Pain 2022;23:754–771.

[13] Cuevas DCS, Ruiz REC, Collina DD, Criollo CJT. Effective brain connectivity related to non-painful thermal stimuli using EEG. Biomed Phys Eng Express 2024;10:045044.

[14] Friston K. A theory of cortical responses. Philos Trans R Soc Lond B Biol Sci 2005;360:815–836.

[15] Gramfort A, Luessi M, Larson E, Engemann DA, Strohmeier D, Brodbeck C, Goj R, Jas M, Brooks T, Parkkonen L, Hämäläinen M. MEG and EEG data analysis with MNE-Python. Front Neurosci 2013;7. doi:10.3389/fnins.2013.00267.

[16] Greffrath W, Baumgärtner U, Treede R-D. Peripheral and central components of habituation of heat pain perception and evoked potentials in humans. PAIN 2007;132:301.

[17] Khona M, Fiete IR. Attractor and integrator networks in the brain. Nat Rev Neurosci 2022;23:744– 766.

[18] Larson E, Gramfort A, Engemann DA, Leppakangas J, Brodbeck C, Jas M, Brooks TL, Sassenhagen J, McCloy D, Luessi M, King J-R, Höchenberger R, Goj R, Brunner C, Favelier G, van Vliet M, Wronkiewicz M, Rockhill A, Holdgraf C, Scheltienne M, Massich J, Appelhoff S, Bekhti Y, Leggitt A, Dykstra A, Trachel R, Luke R, De Santis L, Panda A, Magnuski M, Westner B, Wakeman DG, Strohmeier D, Bharadwaj H, Linzen T, Barachant A, Ruzich E, Bailey CJ, Li A, Moutard C, Bloy L, Raimondo F, Nurminen J, Billinger M, Montoya J, Woodman M, Huberty S, Lee I, Schulz M, Foti N, Nangini C, García Alanis JC, Orfanos DP, Hauk O, Maddox R, LaPlante R, Drew A, Dinh C, Dumas G, Martin, Benerradi J, Hartmann T, Ort E, Billinger M, Pasler P, Repplinger S, Rudiuk A, Radanovic A, Buran B, Woessner J, Massias M, Hämäläinen M, Sripad P, Chirkov V, Mullins C, Raimundo F, Kaneda M, Alday P, Pari R, Kornblith S, Halchenko Y, Luo Y-H, Gramfort A, Kasper J, Doelling K, Jensen M, Gahlot T, Binns TS, Nunes A, Gütlin D, Heinila E, Armeni K, kjs, Weinstein A, Lamus C, Galván CM, Moënne-Loccoz C, Altukhov D, Peterson E, Hanna J, Houck J, Klein N, Roujansky P, Luke R, Ruuskanen S, Kern S, Rantala A, Maess B, Forster C, O’Reilly C, Welke D, Kolkhorst H, Banville H, Zhang J, Maksymenko K, Clarke M, Anelli M, Chapochnikov N, Bannier P-A, Choudhary S, Kim C, Klotzsche F, Wong F-T, Kojcic I, Nielsen JD, Lankinen K, Tabavi K, Thibault L, Gerster M, Alibou N, Gayraud N, Ward N, Herbst S, Férat V, Radanovic A, Quinn A, Gauthier A, Pinsard B, Welke D, Stephen E, Hornberger E, Hathaway E, Kalenkovich E, Mamashli F, Belonosov G, O’Neill G, Marinato G, Anevar H, Abdelhedi H, Sosulski J, Stout J, Calder-Travis J, Zhu JD, Eisenman L, Esch L, Dovgialo M, Barascud N, Legrand N, Kapralov N, Chu Q, Falach R, Deslauriers-Gauthier S, Cotroneo S, Matindi S, Bierer S, Binns TS, Stenner T, Peterson V, Baratz Z, Tonin A, Kovrig A, Pascarella A, Karekal A, de la Torre C, Gohil C, Zhao C, Krzemiński D, Makowski D, Mikulan E, Hofer F, Schiratti J-B, Evans J, Veillette J, Drew J, Teves J, Mathewson K, Gwilliams L, Varghese L, Hamilton L, Gemein L, Hecker L, Lx37, van Es M, Boggess M, Eberlein M, Žák M, Sherif M, Kozhemiako N, Srinivasan N, Wilming N, Kozynets O, Molfese PJ, Ablin P, Das P, Bertrand Q, Shoorangiz R, Scholz R, Hübner R, Sommariva S, Er S, Khan S, Datta S, Papadopoulo T, Donoghue T, Jochmann T, Merk T, Flak T, Dupré la Tour T, NessAiver T, akshay0724, sviter, Earle-Richardson A, Hindle A, Koutsou A, Fecker A, Wagner A, Ciok A, Lepauvre A, Kiefer A, Gilbert A, Pradhan A, Padee A, Dubarry A-S, Waniek AN, Singhal A, Rokem A, Pelzer A, Hurst A, Beasley B, Nicenboim B, de la Torre C, Clauss C, Mista C, Li C-H, Braboszcz C, Schad DC, Hasegan D, Tse D, Sleiter DE, Haslacher D, Sabbagh D, Kostas D, Petkova D, Issagaliyeva D, Das D, Wetzel D, Eich E, DuPre E, Lau E, Olivetti E, Varano E, Altamiranda E, Brayet E, de Montalivet E, Goldstein E, Mamashli F, Negahbani F, Zamberlan F, Pop F, Weber FD, Tan G, Brookshire G, O’Neill G, Giulio, Reina G, Maymandi H, Arzoo HA, Sonntag H, Ye H, Shin H, Elmas HO, Azz I, Machairas I, Zubarev I, de Jong I, Phelan J, Kaczmarzyk J, Zerfowski J, van den Bosch JJF, Van Der Donckt J, van der Meer J, Niediek J, Koen J, Bear JJ, Dammers J, Galán JGN, Welzel J, Slama K, Leinweber K, Grabot L, Andersen LM, Almeida LR, Barbosa LS, Alfine L, Hejtmánek L, Balatsko M, Kitzbichler M, Kumar M, Kadwani M, Sutela M, Koculak M, Henney M, BaBer M, Oberg M, van Harmelen M, Courtemanche M, Tucker M, Visconti di Oleggio Castello M, Dold M, Toivonen M, Shader M, Cespedes M, Krause M, Rybář M, He M, Daneshzand M, Fourcaud-Trocmé N, Gensollen N, Proulx N, Focke N, Chalas N, Markowitz N, Shubi O, Mainar P, Sundaram P, Silva P, Li Q, Barthélemy Q, Nadkarni R, Gatti R, Apariciogarcia R, Aagaard R, Nasri R, Koehler R, Stargardsky R, Oostenveld R, Seymour R, Schirrmeister RT, Law R, Pai S, Perry S, Louviot S, Saha S, Mathot S, Major S, Treguer S, Castaño S, Deng S, Antopolskiy S, Shirazi S (Yahya), Wong S, Wong S, Hofmann SM, Poil S-S, Foslien S, Singh S, Chambon S, Bethard S, Gutstein SM, Meyer SM, Wang T, Moreau T, Radman T, Gates T, Ma T, Stone T, Clausner T, Anijärv TE, Kumaravel VP, Turner W, Zuazo X de, Xia X, Zuo Y, Zhang Z, Zeng Z, btkcodedev, buildqa, luzpaz. MNE-Python. 2024. doi:10.5281/zenodo.14519545.

[19] Marshall W, Gomez-Ramirez J, Tononi G. Integrated Information and State Differentiation. Front Psychol 2016;7. doi:10.3389/fpsyg.2016.00926.

[20] Mulders D, Seymour B, Mouraux A, Mancini F. Confidence of probabilistic predictions modulates the cortical response to pain. Proc Natl Acad Sci 2023;120:e2212252120.

[21] Nuñez-Ibero M, Camino-Pontes B, Diez I, Erramuzpe A, Martinez-Gutierrez E, Stramaglia S, Alvarez-Cienfuegos JO, Cortes JM. A Controlled Thermoalgesic Stimulation Device for Exploring Novel Pain Perception Biomarkers. IEEE J Biomed Health Inform 2021;25:2948–2957.

[22] Orlowski P, Bola M. Sensory modality defines the relation between EEG Lempel–Ziv diversity and meaningfulness of a stimulus. Sci Rep 2023;13:3453.

[23] R: The R Project for Statistical Computing. 2025. Available: https://www.r-project.org/. Accessed 4 Jul 2025.

[24] Ronga I, Valentini E, Mouraux A, Iannetti GD. Novelty is not enough: laser-evoked potentials are determined by stimulus saliency, not absolute novelty. J Neurophysiol 2013;109:692–701.

[25] Sarasso S, Casali AG, Casarotto S, Rosanova M, Sinigaglia C, Massimini M. Consciousness and complexity: a consilience of evidence. Neurosci Conscious 2021;2021:niab023.

[26] Sinitsyn DO, Poydasheva AG, Bakulin IS, Legostaeva LA, Iazeva EG, Sergeev DV, Sergeeva AN, Kremneva EI, Morozova SN, Lagoda DY, Casarotto S, Comanducci A, Ryabinkina YV, Suponeva NA, Piradov MA. Detecting the Potential for Consciousness in Unresponsive Patients Using the Perturbational Complexity Index. Brain Sci 2020;10:917.

[27] Tayeb Z, Dragomir A, Lee JH, Abbasi NI, Dean E, Bandla A, Bose R, Sundar R, Bezerianos A, Thakor NV, Cheng G. Distinct spatio-temporal and spectral brain patterns for different thermal stimuli perception. Sci Rep 2022;12:919.

[28] Tononi G. An information integration theory of consciousness. BMC Neurosci 2004;5:42.

[29] Torta DM, Liang M, Valentini E, Mouraux A, Iannetti GD. Dishabituation of laser-evoked EEG responses: dissecting the effect of certain and uncertain changes in stimulus spatial location. Exp Brain Res 2012;218:361–372.

[30] Vignesh D, He S, Banerjee S. A review on the complexities of brain activity: insights from nonlinear dynamics in neuroscience. Nonlinear Dyn 2025;113:4531–4552.

[31] Wang Y, Niu Z, Xia X, Bai Y, Liang Z, He J, Li X. Application of Fast Perturbational Complexity Index to the Diagnosis and Prognosis for Disorders of Consciousness. IEEE Trans Neural Syst Rehabil Eng 2022;30:509–518.

[32] Watanabe H, Shibuya S, Masuda Y, Sugi T, Saito K, Nagashima K. Spatial and temporal patterns of brain neural activity mediating human thermal sensations. Neuroscience 2025;564:260–270.

[33] Wei M, Liao Y, Liu J, Li L, Huang G, Huang J, Li D, Xiao L, Zhang Z. EEG Beta-Band Spectral Entropy Can Predict the Effect of Drug Treatment on Pain in Patients With Herpes Zoster. J Clin Neurophysiol 2022;39:166.

[34] Wu D, Cai G, Yuan Y, Liu L, Li G, Song W, Wang M. Application of nonlinear dynamics analysis in assessing unconsciousness: A preliminary study. Clin Neurophysiol 2011;122:490–498.

