## Supplementary figures and tables for "Perturbational complexity of cortical responses to thermal pain"

### Supplementary material

**Table S1. Model 1, ANOVA**

|  | **Sum of squares** | **df** | **statistic** | ***p*** |
| --- | --- | --- | --- | --- |
| **Intercept** | 163.13 | 1 | 174.47 | **< .001** |
| **Temperature** | 31.01 | 2 | 16.58 | **< .001** |
| **Agonist** | 8.63 | 3 | 3.08 | **.028** |
| **Temperature:Agonist** | 6.95 | 6 | 1.24 | .286 |
| **Temperature:Agonist:Patch** | 25.12 | 12 | 2.24 | **.010** |
| **Residuals** | 299.19 | 320 |  |  |

**Table S2. Model 1, post-hoc comparisons between temperatures**

| **Contrast** | **Estimate (Δ)** | **SE** | **df** | **t** | ***p*** | ***p_Holm-Bonferroni_*** |
| --- | --- | --- | --- | --- | --- | --- |
| 42°C - 10°C | -1.43 | 0.13 | 320 | -1.73 | -1.13 | **< .001** |
| 42°C - 60°C | -1.21 | 0.13 | 320 | -1.52 | -0.89 | **< .001** |
| 10 °C - 60 °C | 0.22 | 0.13 | 320 | -0.09 | 0.53 | .0852 |

**Table S3. Model 1, post-hoc comparisons for different temperatures and agonists**

Contrasts are calculated within patch conditions (vehicle vs agonist). Holm bonferroni correction has been applied inside each Agonist group to control the family-wise type I error.

| **Temperature** | **Agonist** | **Δ_vehicle - agonist_** | **SE** | **df** | **t** | ***p*** | ***p_Holm-Bonferroni_*** |
| --- | --- | --- | --- | --- | --- | --- | --- |
| 42 | Capsaicin (0.25%) | -0.27 | 0.36 | 320 | -0.98 | 0.453 | 0.906 |
| 10 | Capsaicin (0.25%) | 0.27 | 0.35 | 320 | -0.42 | 0.441 | 0.883 |
| 60 | Capsaicin (0.25%) | 0.18 | 0.37 | 320 | -0.55 | 0.621 | 1.000 |
| 42 | Capsaicin (1%) | 0.03 | 0.36 | 320 | -0.68 | 0.942 | 0.942 |
| 10 | Capsaicin (1%) | 0.91 | 0.37 | 320 | 0.19 | 0.013 | **0.040** |
| 60 | Capsaicin (1%) | 0.21 | 0.38 | 320 | -0.54 | 0.589 | 1.000 |
| 42 | Cinnamic aldehyde (10%) | 0.37 | 0.34 | 320 | -0.3 | 0.277 | 0.867 |
| 10 | Cinnamic aldehyde (10%) | 0.08 | 0.34 | 320 | -0.59 | 0.816 | 0.883 |
| 60 | Cinnamic aldehyde (10%) | 0.59 | 0.35 | 320 | -0.1 | 0.096 | 0.382 |
| 42 | Menthol (20%) | 0.47 | 0.38 | 320 | -0.28 | 0.217 | 0.867 |
| 10 | Menthol (20%) | 1.23 | 0.35 | 320 | 0.54 | <0.001 | **0.002** |
| 60 | Menthol (20%) | 0.38 | 0.39 | 320 | -0.4 | 0.336 | 1.000 |

**Table S4. Model 2, Coefficients**

|  | **Estimate (β)** | **SE** | **t** | ***p*** |
| --- | --- | --- | --- | --- |
| Intercept | 5.32 | 0.07 | 80.77 | **< .001** |
| Temperature I2 | -0.35 | 0.09 | -3.70 | **< .001** |
| p(I1) | 0.14 | 0.07 | 2.12 | **.034** |
| Temperature I2 x p(I1) | -0.24 | 0.09 | -2.55 | **.011** |

**Table S5. Model 2, Average marginal effects**

|  | **AME** | **SE** | **z** | ***p*** |
| --- | --- | --- | --- | --- |
| p(I1) | 0.02 | 0.05 | 0.45 | .655 |
| Temperature I2 | -0.35 | 0.09 | -3.71 | **< .001** |

**Table S6. Model 2, Marginal effects of p(I1) per temperature**

| **Temperature** | **ME** | **SE** | **df** | ***p*** |
| --- | --- | --- | --- | --- |
| I1 | 0.14 | 0.07 | 591 | **.034** |
| I2 | -0.10 | 0.07 | 591 | .137 |

**Table S7. Model 3, coefficients**

|  | **Estimate (β)** | **SE** | **t** | ***p*** |
| --- | --- | --- | --- | --- |
| Intercept | 6.51 | 0.49 | 13.22 | **< .001** |
| Type (hot) | -5.68 | 0.72 | -7.89 | **< .001** |
| Temperature | -0.10 | 0.04 | -2.47 | **.014** |
| Type (hot) : Temperature | 0.17 | 0.04 | 4.17 | **< .001** |

**Table S8. Model 3, marginal effects of temperature**

| **Type** | **ME** | **SE** | **df** | ***p*** |
| --- | --- | --- | --- | --- |
| cold | -0.10 | 0.04 | 295 | **.014** |
| hot | 0.07 | 0.01 | 295 | **< .001** |

**Table S9**. **Regression coefficients for Dataset 1, cold**

|  | **Estimate (β)** | **SE** | **t** | ***p*** |
| --- | --- | --- | --- | --- |
| **Intercept** | 0.11 | 0.10 | 1.12 | .264 |
| N2-P2 Amplitude | 0.20 | 0.11 | 1.89 | .062 |
| N2 Latency | -0.17 | 0.11 | -1.48 | .143 |
| P2 Latency | 0.10 | 0.12 | 0.83 | .409 |
| N2 P2 Amplitude:N2 Latency | 0.08 | 0.13 | 0.63 | .530 |
| N2-P2 Amplitude:P2 Latency | -0.01 | 0.17 | -0.03 | .973 |
| N2 Latency:P2 Latency | 0.01 | 0.10 | 0.08 | .937 |
| N2-P2 Amplitude:N2 Latency:P2 Latency | 0.07 | 0.14 | 0.49 | .625 |

**Table S10**. **Marginal effects for Dataset 1 Model 4, cold**

|  | **AME** | **SE** | **z** | ***p*** |
| --- | --- | --- | --- | --- |
| N2 Latency | -0.17 | 0.11 | -1.54 | .124 |
| N2-P2 Amplitude | 0.23 | 0.10 | 2.25 | **.024** |
| P2 Latency | 0.09 | 0.12 | 0.76 | .448 |

**Table S11**. **Regression coefficients for Dataset 1 Model 4, hot**

|  | **Estimate (β)** | **SE** | **t** | ***p*** |
| --- | --- | --- | --- | --- |
| Intercept | 0.12 | 0.12 | 1.01 | .316 |
| N2-P2 Amplitude | -0.03 | 0.12 | -0.23 | .819 |
| N2 Latency | -0.24 | 0.13 | -1.91 | .060 |
| P2 Latency | -0.11 | 0.12 | -0.93 | .356 |
| N2P2 Amplitude:N2 Latency | -0.14 | 0.13 | -1.04 | .299 |
| N2-P2 Amplitude:P2 Latency | 0.04 | 0.14 | 0.3 | .763 |
| N2 Latency:P2 Latency | -0.02 | 0.14 | -0.12 | .904 |
| N2-P2 Amplitude:N2 Latency:P2 Latency | 0.15 | 0.14 | 1.05 | .296 |

**Table S12**. **Marginal effects for Dataset 1 Model 4, hot**

|  | **AME** | **SE** | **z** | ***p*** |
| --- | --- | --- | --- | --- |
| N2 Latency | -.25 | 0.13 | -1.94 | .052 |
| N2-P2 Amplitude | 0.02 | 0.11 | 0.21 | .835 |
| P2 Latency | -0.14 | 0.11 | -1.32 | .188 |

**Table S13**. **Regression coefficients for Dataset 2 Model 4, cold**

|  | **Estimate (β)** | **SE** | **t** | ***p*** |
| --- | --- | --- | --- | --- |
| Intercept | 0.02 | 0.05 | 0.46 | .647 |
| N2-P2 Amplitude | 0.02 | 0.05 | 0.32 | .750 |
| N2 Latency | 0.03 | 0.05 | 0.48 | .630 |
| P2 Latency | -0.02 | 0.05 | -0.45 | .655 |
| N2P2 Amplitude:N2 Latency | 0.15 | 0.07 | 2.24 | **.025** |
| N2-P2 Amplitude:P2 Latency | -0.10 | 0.07 | -1.35 | .176 |
| N2 Latency:P2 Latency | 0.04 | 0.04 | 1.01 | .312 |
| N2-P2 Amplitude:N2 Latency:P2 Latency | 0.04 | 0.08 | 0.49 | .626 |

**Table S14**. **Marginal effects for Dataset 2 Model 4, cold**

|  | **AME** | **SE** | **z** | ***p*** |
| --- | --- | --- | --- | --- |
| N2 Latency | 0.03 | 0.05 | 0.67 | .503 |
| N2-P2 Amplitude | 0.03 | 0.06 | 0.57 | .570 |
| P2 Latency | -0.02 | 0.05 | -0.32 | .752 |

**Table S15**. **Regression coefficients for Dataset 2 Model 4, hot**

|  | **Estimate (β)** | **SE** | **t** | ***p*** |
| --- | --- | --- | --- | --- |
| Intercept | 0.20 | 0.05 | 4.26 | **< .001** |
| N2-P2 Amplitude | 0.01 | 0.05 | 0.19 | .848 |
| N2 Latency | 0.21 | 0.05 | 4.14 | **< .001** |
| P2 Latency | -0.05 | 0.05 | -0.99 | .323 |
| N2P2 Amplitude:N2 Latency | < 0.01 | 0.05 | 0.04 | .971 |
| N2-P2 Amplitude:P2 Latency | 0.08 | 0.06 | 1.31 | .190 |
| N2 Latency:P2 Latency | -0.05 | 0.05 | -0.91 | .362 |
| N2-P2 Amplitude:N2 Latency:P2 Latency | 0.11 | 0.06 | 1.98 | **.049** |

**Table S16**. **Marginal effects for Dataset 2 Model 4, hot**

|  | **AME** | **SE** | **z** | ***p*** |
| --- | --- | --- | --- | --- |
| N2 Latency | 0.20 | 0.05 | 4.02 | **< .001** |
| N2-P2 Amplitude | 0.04 | 0.05 | 0.68 | .496 |
| P2 Latency | -0.03 | 0.05 | -0.64 | .523 |

**Table S17. R^2^ for Model 4 applied to different dataset**

| **Regression** | **R^2^** | **SE** | **LCL** | **UCL** |
| --- | --- | --- | --- | --- |
| Dataset 1, cold | .10 | 0.16 | -.22 | .42 |
| Dataset 1, hot | .10 | 0.14 | -.17 | .38 |
| Dataset 2, cold | .02 | 0.10 | -.17 | .21 |
| Dataset 2, hot | .05 | 0.15 | -.23 | .34 |

**Supplementary analyses - Dependency of PCI from numbers of epochs averaged**

To verify the stability of PCI index when averaging the ERPs for different numbers of epochs, we selected the conditions that had at least 51 good epochs ($p$(stimulus) = 0.7) in dataset 2. The following model was implemented:

PCI ~ Temperature*epochs_number

Where Temperature is a categorical variable, and epochs_number a mean-centered metric variable. Regression coefficients did not show any significant effect of the number of epochs on PCI values, for both temperatures, but confirmed higher PCI values for cold stimuli compared to hot ones (Table 18)

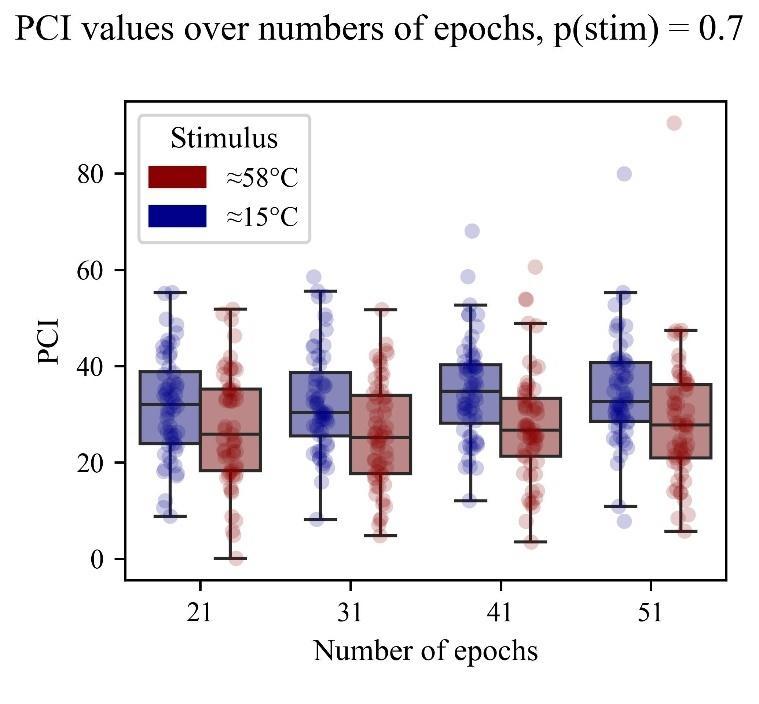

**Figure S1. PCI dependence on the number of averaged epochs**

The box‑and‑whisker plots show raw PCI scores grouped by stimulus (I1≈ 15 °C, blue; I2≈ 58 °C, red. Each semi‑transparent dot represents the single PCI estimate for one participant; boxes mark the median and IQR and whiskers extend to 1.5 × IQR. Consistent with the statistical analysis (Table 18) it is not possible to observe a significant effect of the number of epochs on PCI for both stimuli, indicating that, at least from 21 epochs onwards, the index is stable across the possible variations of SNR that could occur by averaging different number of trials.

**Table S18. Number of epochs effect on PCI**

|  | **Estimate (β)** | **SE** | **t** | ***p*** |
| --- | --- | --- | --- | --- |
| Intercept | 33.38 | 0.69 | 48,11 | **< .001** |
| Number of epochs | 0.06 | 0.04 | 1.34 | **.181** |
| Temperature (hot) | -6.35 | 0.99 | -6.44 | **< .001** |

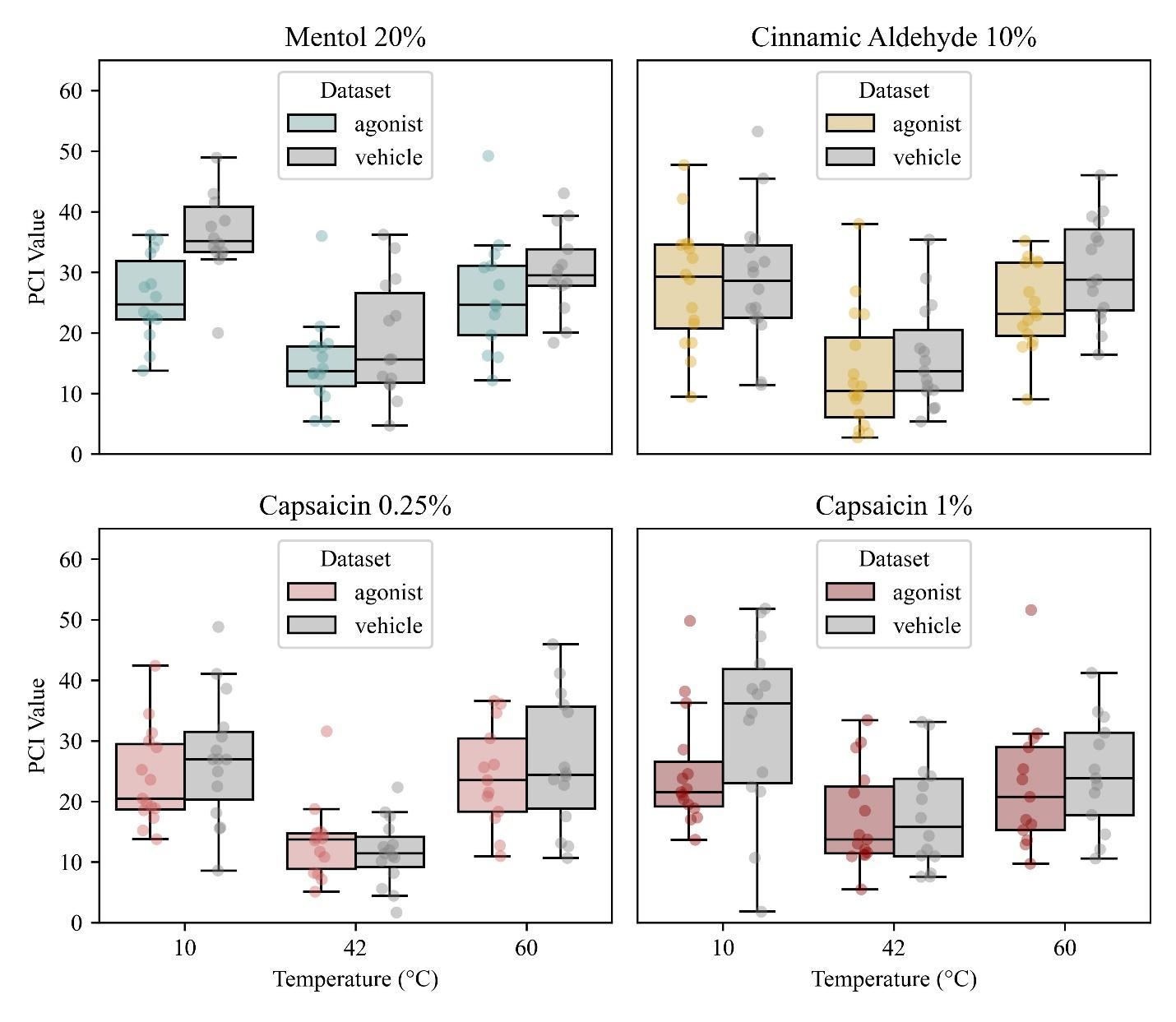

**Figure S2. PCI values as a function of temperature and topical patch in Dataset 1.**

Box‑and‑whisker plots show raw PCI scores (32‑channel montage) for all the agonists employed in Dataset 1. For each agonist, participants received a vehicle patch and an active patch before stimulation at 10 °C, 42 °C, and 60 °C; the patch order was counter‑balanced within subjects. Semi‑transparent dots represent the single PCI estimate for every participant‑temperature‑patch combination, boxes mark the median and IQR, and whiskers extend to 1.5 × IQR.

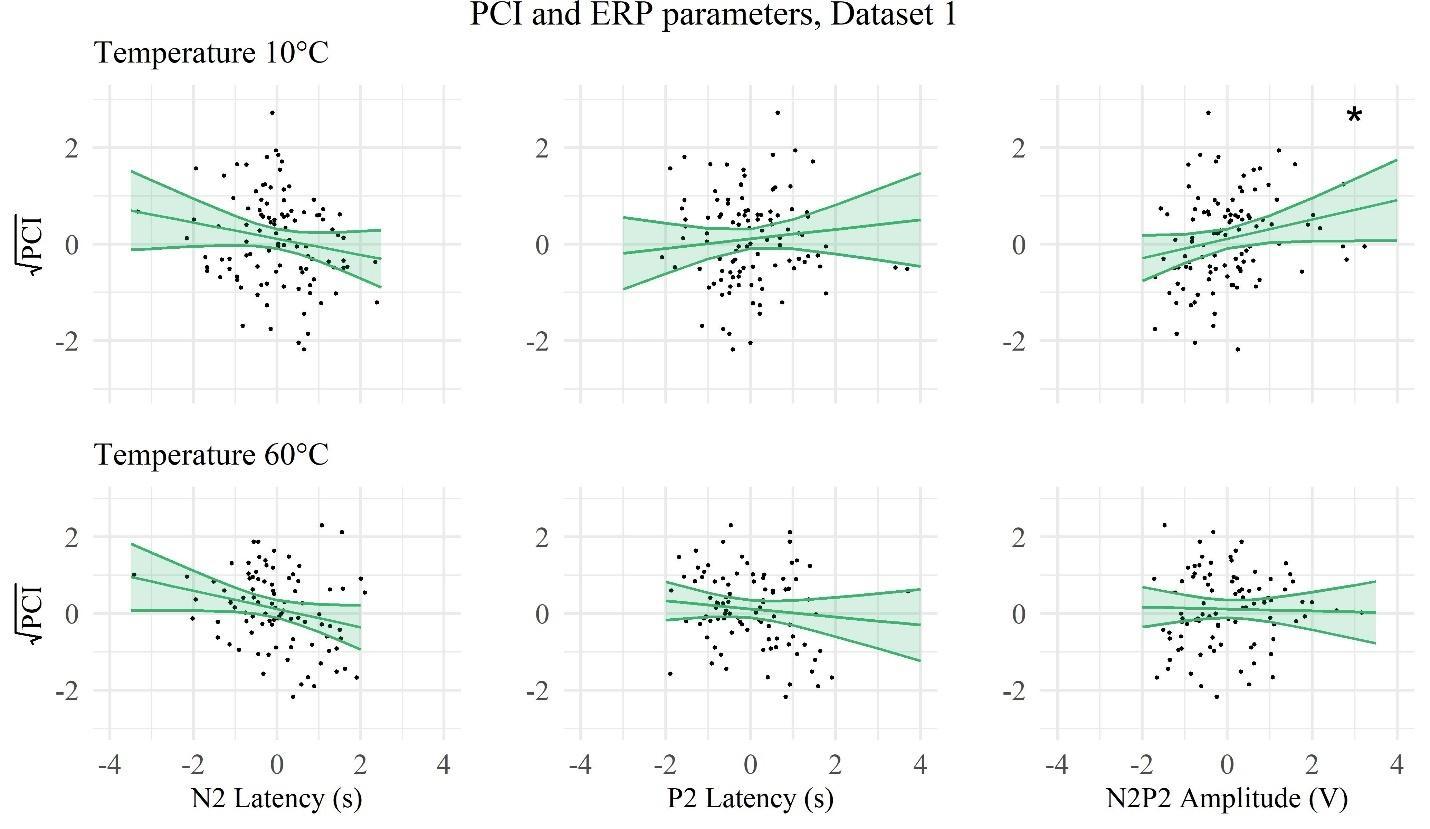

**Figure S3. Relationship between PCI and N2–P2 peak parameters in Dataset 1.**

Each panel shows z‑scored PCI plotted against N2 latency, P2 latency, or N2–P2 amplitude; the upper row corresponds to 10 °C, the lower row to 60 °C. Solid lines are least‑squares fits with 95 % confidence bands. For cold temperature, N2-P2 Amplitude had a significant effect on PCI (AME = .23, p = .024, significance indicated in the plot by the symbol *), while no other significant effect was found between peak parameters and complexity. All the variables in the plot have been mean-scaled and centered.

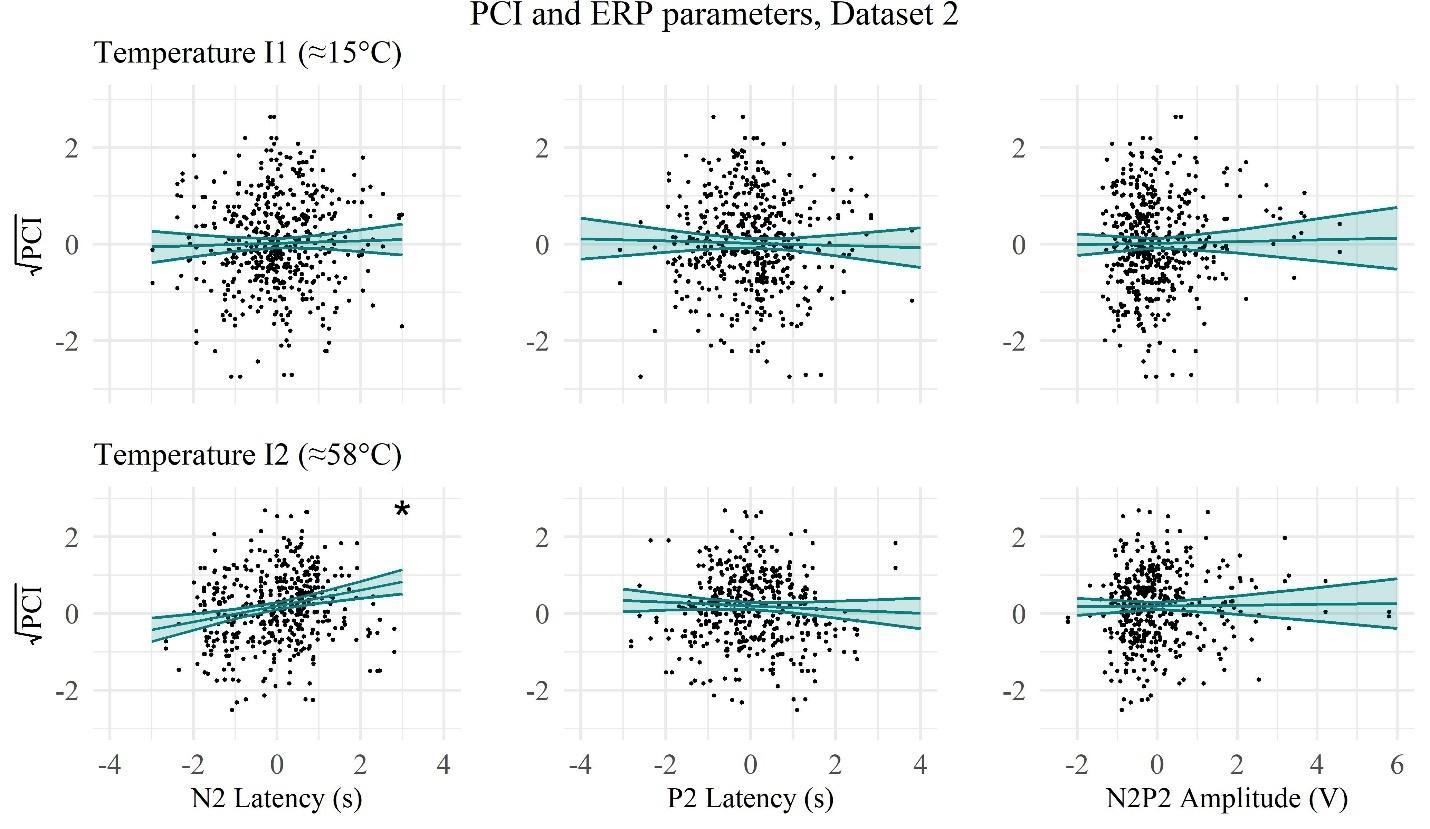

**Figure S4. Relationship between PCI and N2–P2 peak parameters in Dataset 2.**

Each panel shows z‑scored PCI plotted against N2 latency, P2 latency, or N2–P2 amplitude; the upper row corresponds to I1≈ 15 °C, the lower row to I2≈ 58°C. Solid lines are least‑squares fits with 95 % confidence bands. For hot stimuli, there was a significant marginal effect of N2 Latency (AME = .20, p < .001, significance indicated in the plot by the symbol *). All the variables have been mean-scaled and centered.
